# Spatial patterns of vascular plant species richness in Poland: relations among species group richness and hot spot locations

**DOI:** 10.1101/2024.07.14.603435

**Authors:** T.H. Szymura, H. Tegegne, M. Szymura

## Abstract

Knowledge of spatial patterns of species richness (SR) is highly relevant for theoretical research in ecology and the development of conservation plans. In Poland, despite a long tradition of botanical surveys, vascular plant SR has not been mapped, nor have the correlations in richness among different plant species groups been explored. Here we used a recently published data set to examine spatial patterns and relationships among the joined SR of vascular plant species, including native species, archeophytes, neophytes, and species with high conservation value (i.e., red list species). The basic spatial unit employed was a 10 × 10 km grid from the Atlas of Distribution of Vascular Plants in Poland (ATPOL). We found that the richness patterns of native species, archeophytes, neophytes, and red-list species were positively correlated. The main patterns of SR and the percentage of particular groups in the joined SR were based on three components: (1) gradient of overall SR, (2) invasion level, and (3) peculiarity of flora in some regions resulting from the high number and proportion of rare species that often have high conservation value. In general, northeastern Poland was species-poor, while the Carpathian Mountain range, the uplands in southern Poland, and some parts of Wisła River valley had the highest SR concentrations. The location of SR hotspots usually did not overlap with the existing system of national parks. The correlations among native SR, high conservation value species, and neophyte SR suggest that biological invasions are among the most important threats to vascular plant diversity in Poland. Finally, we demonstrated that the presented maps, despite likely biases in SR assessments, seem to reflect general ecological gradients influencing vascular plant distribution in Poland.

## 1. Introduction

Plants constitute most of the biomass of terrestrial ecosystems (Bar-On et al., 2018), and the species richness (SR) of vascular plants can serve to predict multi-taxon SR (Brunbjerg, 2018). This latter information helps in site selection for conservation of numerous other groups of organisms (e.g., spiders, carbides, bryophytes; Sætersdal et al., 2004). Unfortunately, perceptual factors such as the lack of motion by plants and their tendency to visually blend into the landscape, along with cultural factors such as a greater focus on animals in formal biological education, lead to “plant blindness,” defined as the tendency to simply overlook plants. As a result, plant conservation initiatives lag and receive considerably less funding than animal conservation projects (Balding & Williams, 2016; Adamo et al., 2022).

The human impact of the Anthropocene has caused unprecedented biodiversity loss (Díaz et al., 2019). Owing to the alarmingly high rates of plant invasions and the range shifts and extinctions of species, documentation of the Earth’s flora has become urgently needed. Comprehensive syntheses of information about individual plant groups underlie a broad range of research, conservation of plant diversity, and science outreach activities to improve awareness of plants (for review see: Grace et al., 2021). However, creating an inventory of biodiversity is challenging and requires significant resources, making the efficient and optimal design of survey and inventory efforts particularly important (Nuñez-Penichet et al., 2022). As a result, only a few countries, such as the Czech Republic (Klinkovská et al., 2024), Denmark (Finderup Nielsen, 2019), Germany (Eichenberg et al., 2021; Jandt et al., 2022), and the United Kingdom (Pescott et al., 2019, Stroh et al., 2023) have developed databases that enable tracing country-scale changes in vascular plants species distribution through past decades. The results of such analyses have confirmed the decline of SR and/or distribution ranges of native species, including rare and protected species, and the simultaneous increases of SR and/or distribution ranges of neophytes in Central Europe (Klinkovská et al., 2024; Eichenberg et al.; 2021; but for another trend in Great Britain, see Montràs-Janer et al., 2022). Moreover, the effects of recent anthropogenic activities are likely more severe than currently known because of the temporal delay in the extinction of numerous species, which will only become apparent in coming decades (Dullinger et al., 2013).

Poland is a large country (terrestrial area ca 312,000 km^2^) that lies mainly in a continental biographic region. The mountains in the south are considered an alpine biogeographical region, while northeastern Poland border with the boreal region (Cervellini et al., 2020). According to the national checklist, the plant SR is assessed at 2,600 native species and 534 neophytes (Mirek et al., 2020). These numbers are comparable to the SR in the Czech Republic (Danihelka et al., 2012), which has about one-fourth the area of Poland. Similar to the flora of other Central European countries, the number of neophytes in Poland has been increasing recently (Jackowiak, 2023). The majority of Poland’s territory is large flat lowlands (**Fig 1a**), which allows easy plant migration from east to west and vice versa (Szafer, 1955). As a result, species from different geographical ranges can be found in Poland (e.g., maps shown by Finnie et al., 2007), while altitudinal differentiation is considered among the most important factors shaping plant species distribution (Szafer, 1955). The land relief and feasible plant migration result in a low number of endemics in Poland. Mirek and Pięknoś-Mirkowa (2009) recorded 169 species and subspecies, including numerous apomictic, so-called ‘microspecies’ from *Hieracium*, *Alchemilla*, and *Taraxacum* genera, whose distribution is mostly restricted to mountains. The Polish flora has a long history of investigation, the general description of flora and vegetation was summarized as early as 1959 (Szafer, 1959) and translated to English a few years later (Szafer, 1966). The Atlas of Distribution of Vascular Plants in Poland (ATPOL) project has been ongoing since the 1970s (Zając, 1978), and the Polish Vegetation Database (PVD) has been operating since 2007 (Kącki & Śliwiński, 2012). Nonetheless, till now there has been no maps of spatial patterns of plant SR for the entire country (but see Moraczewski & Sudnik-Wojciechowska, 2007).

**Figure 1.**
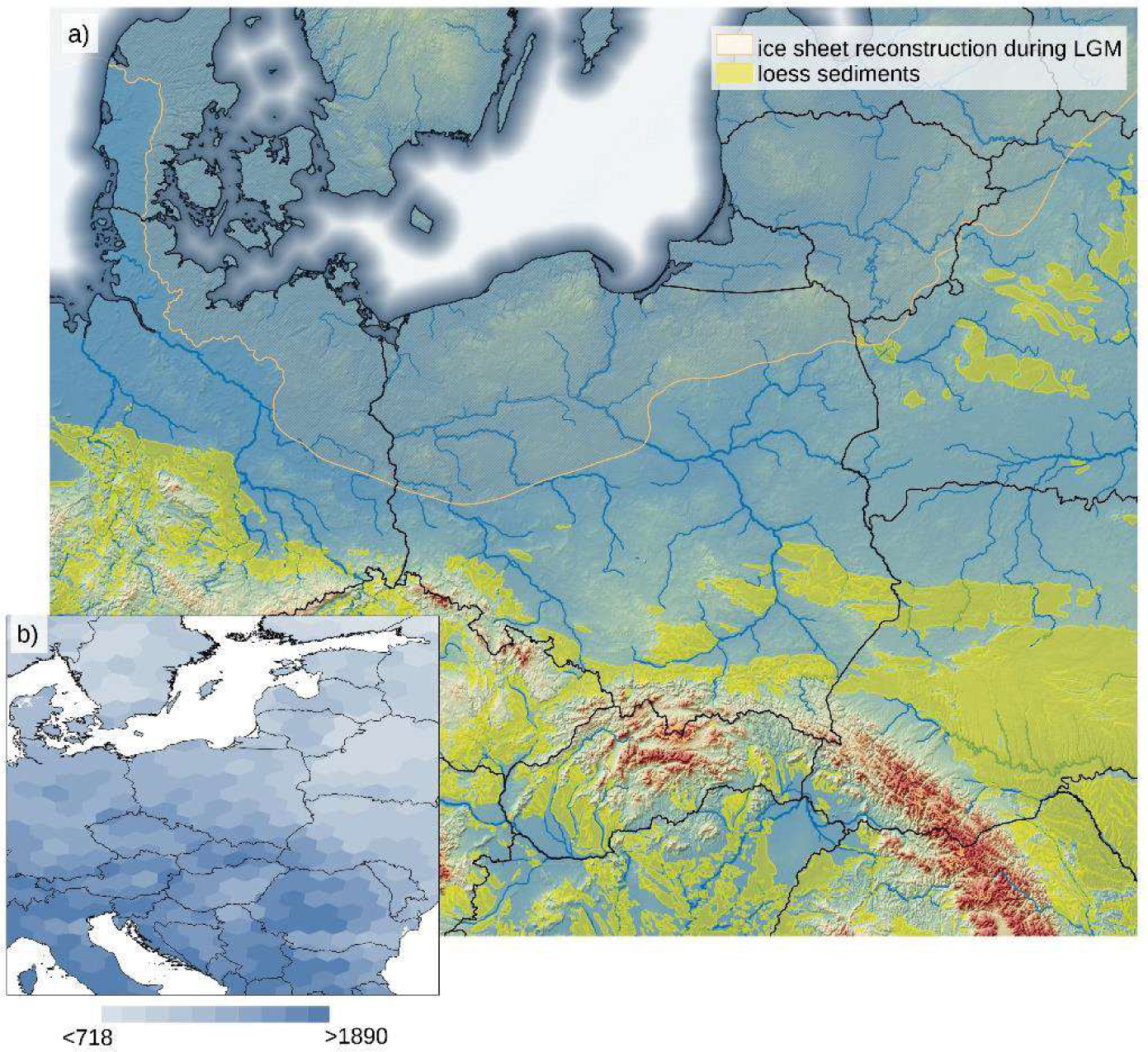
Location of Poland and neighboring countries on the background of land relief, ice sheet cover during Last Glaciation Maximum (LGM), the location of loess sediments, the network of main rivers (a), and the map species richness distribution in Europe (b). The ice sheet reconstruction following EPHA (2019), loess sediments from Lehmkul et al. (2020), river network according to AQUSTAT (2021), and global model of species richness from Cai et al. (2022).

In this article, we determine vascular plant SR patterns on a spatial scale that encompasses the entirety of Poland, using a new comprehensive data set. We explore the location of biodiversity hot spots and cold spots, spatial patterns of invasion, and correlations between particular species groups. The results are then presented on maps.

## 2. Material and Methods

### 2.1. Species richness data set

The data used in this study are from the recently published data set ‘Spatial pattern of vascular plant species richness in Poland,’ available from the Zenado repository (Szymura et al., 2023a) and described in detail by Szymura et al. (2023c). In brief, the data show SR in more than 3000 squares (10 ×10 km) resulting from combining and harmonizing data from the ATPOL (Zając, 1978) and PVD (Kącki & Śliwiński, 2012) projects. Besides the joined SR, the data set divides the species into particular groups regarding their origin (native, neophyte, archeophyte), conservation value (red list species), ecology (apophyte), and frequency of occurrence (common, moderate, and rare). Although the taxa list in the database includes operational taxonomical units at different taxonomical levels (species, species sensu lato, aggregations), they are referred to in this study as ‘species’ for simplification. Additionally, the number of genera and families are shown in a square, as well as the calculated percentage of particular species groups, with 100% being considered the joined SR.

### 2.2. Analysis method

To produce a map of SR, we used only squares for which more than 80% of the area was within the terrestrial territory of Poland. For further calculations (e.g., identifying hot spots), we excluded the squares from this subset that were identified as being undersampled. Ultimately, we analyzed 2137 species in 2866 squares (for details, see Szymura et al., 2023c), and this information was used for descriptive statistics. The number of squares in which a particular species was found was counted to obtain an overall pattern of spatial distribution of the species in Poland. Further, data from the selected squares were standardized and then explored using principal component analysis (PCA). In additional, Spearman’s rank correlations among all the variables were calculated.

The hot spots and cold spots of SR were defined as 5% squares with the highest and the lowest number of species in given group (Prendergast et al., 1993; Reid, 1998). It was around 144 squares per group: these results were obtained by counting the number of species belonging to a particular species group and selecting squares based on the specific number of species results that varied from the number of distinguished squares for individual species groups, depending on the adopted threshold for the number of species. Next, the squares defined as hot or cold spots were spatially aggregated by combining squares touching each other (including corners) into connected areas. The locations of the hot- and cold-spot regions were assigned to the physical-geographical mesoregions of Poland on maps created by Solon et al. (2018). We decided to use this division instead of the geobotanical regionalization of Poland by Matuszkiewicz (2008) because the first maps are publicly available in recent GIS format.

All the calculations were done in R environment R Core Team (2023), while maps were prepared in QGIS (QGIS.org 2023) software.

## 3. Results

The median value of SR per square was 522 species, most of them were native species (median 436 species, 84%) and archeophytes (median 62, 12%). The high conservation value species accounted for approximately 6% of all species, with a median value of their richness per square of 27. Neophytes had a similar median, with 29 species. The five most frequent species in a particular species group are listed in **Table 1**. The rare species were approximately 11% of all species in a square (average 53 species). Meanwhile, the percentages of moderate and common species were 47% and 41%, respectively, corresponding to 256 and 212 species per square on average. The apophytes constituted an average of 19% of all species, yielding a median value of 100 apophyte species in a square. The SR distribution in squares for particular groups is shown in **Fig. S1,** while descriptive statistics are in **Table S1** in the Appendix. Most of the species have a limited range of distribution in Poland, as shown by the number of occupied squares: about 40% of all species were found in less than 100 squares, while the range of distribution of about 50% of the analyzed species was less than 200 squares (**Fig. 2**).

**Fig 2.**
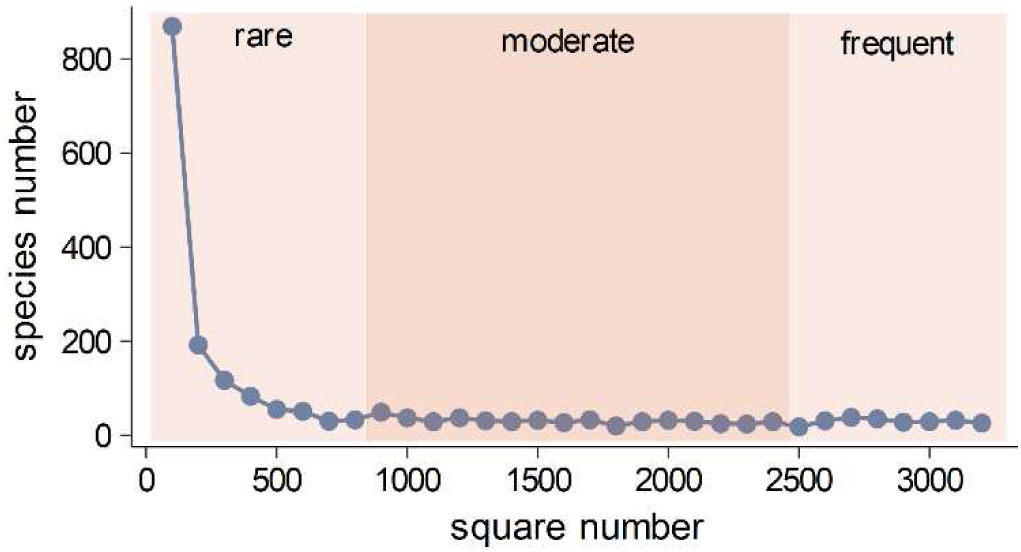
The number of species (y-axis) occurring in a particular square number (x-axis). The different background colors highlight the rare, moderate, and common species groups (see Materials and Methods).

**Table 1.** The median species richness (SR) values of species belonging to a particular group found in Poland, with the five most frequent species in a particular group (species) and the minimum number of squares in which the species had to be detected to be listed in the ‘species’ column

| Species group | Median SR | Five most frequent species | Min. no. of squares |
| --- | --- | --- | --- |
| Native | 437 | *Urtica dioica* s.l., *Achillea millefolium* agg., *Ranunculus acris* agg., *Plantago major* s.l., *Deschampsia cespitosa* | 3173 |
| Red list | 27 | *Aethusa cynapium* s.l., *Agrostemma githago*, *Dactylorhiza majalis* agg., *Lycopodium clavatum*, *Lycopodium annotinum* | 1603 |
| Neophytes | 29 | *Matricaria discoidea*, *Erigeron canadensis*, *Galinsoga parviflora*, *Amaranthus retroflexus*, *Veronica persica* | 2335 |
| Arheophytes | 62 | *Capsella bursa-pastoris*, *Silene latifolia*, *Fallopia convolvulus*, *Viola arvensis*, *Cyanus segetum* | 3008 |

The distribution of SR was not random, and clusters of squares with higher or lower SR were visible. In general, northeastern Poland was poorer in species, while the southern and southeastern areas were species-rich (**Fig 3**). The percentage of rare species was especially high in the mountains, while moderately frequent in uplands and lowlands (**Fig 3e and f**).

**Figure 3.**
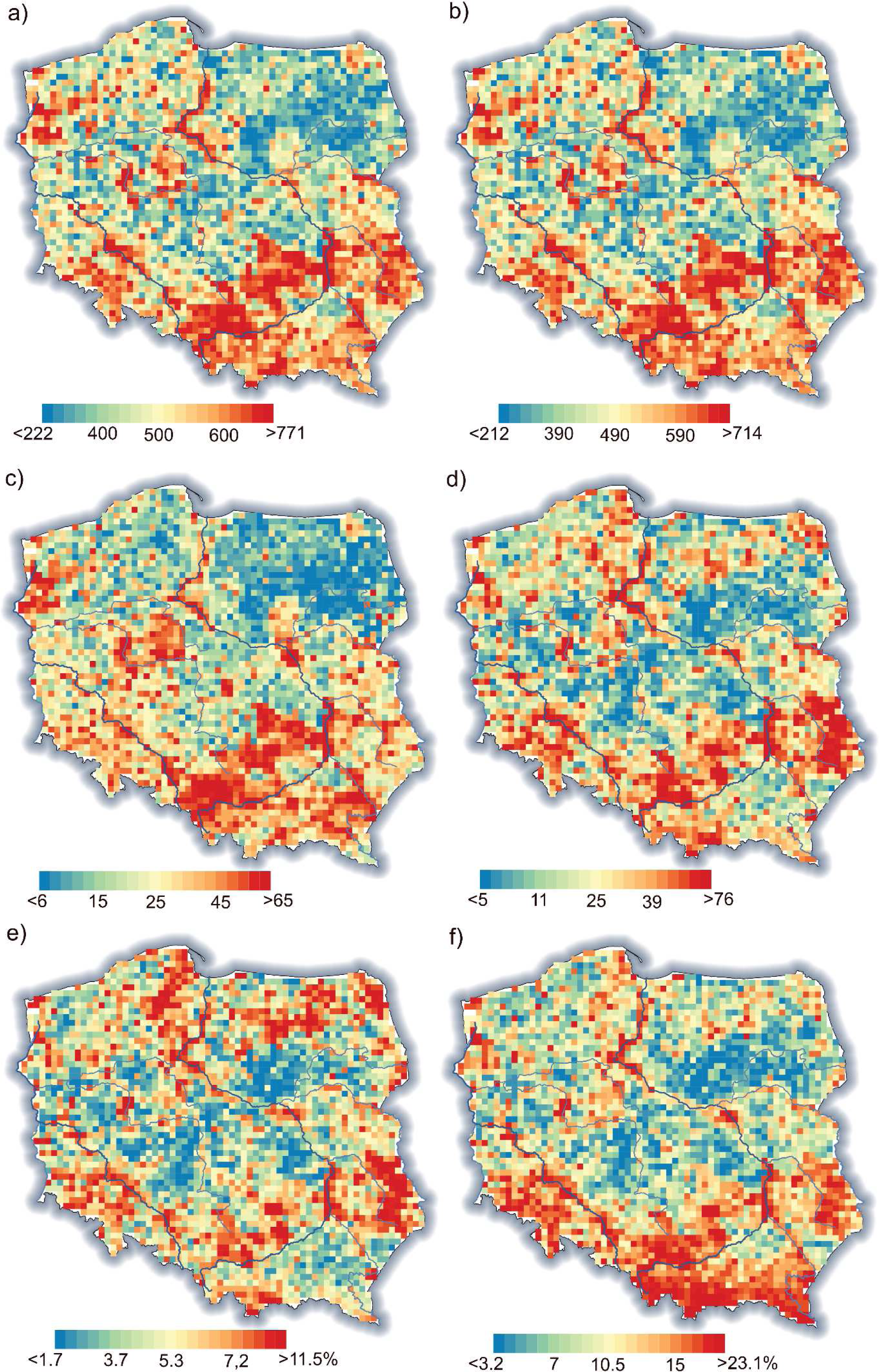
Maps of joined species richness (a), native + archeophytes (b), neophytes (c), and red list (d) species richness, as well as rare (e) and moderate (f) species percentages. On the maps, the undersampled squares are also shown (for details see Methods). The blue lines represent the main watercourses.

Numerous correlations were found between the richness of species in different groups as well as their percentages (**Table S2, Appendix**). In general, the joined SR was positively correlated with SR in all remaining groups and the number of genera and families in a square (**Table S2, Appendix**). The main patterns of SR in a particular group could be attributed to three components, depicted as axes in the PCA results (**Table 2**, **Fig 4**). The first axis (PCA 1) relates to the overall SR gradient, whereby the species-poor squares were dominated by common and moderately frequent species with a high fraction of apophytes, while the species-rich plots were rich in species from all groups as, well family and genera numbers. The species-rich squares also had a high fraction of neophytes, archeophytes, and red-list species. The second axis (PCA 2) underlines the contrast between plots dominated by native species and those with a high fraction of neophytes and archeophytes. The third axis (PCA 3) shows plots with high numbers and percentages of red list species and rare species compared with plots with a high number of common species and a high fraction of moderately frequent species (**Table 2**, **Fig 4**). Thus, PCA 3 can be considered as being associated with a ‘rarity’ gradient.

**Figure 4.**
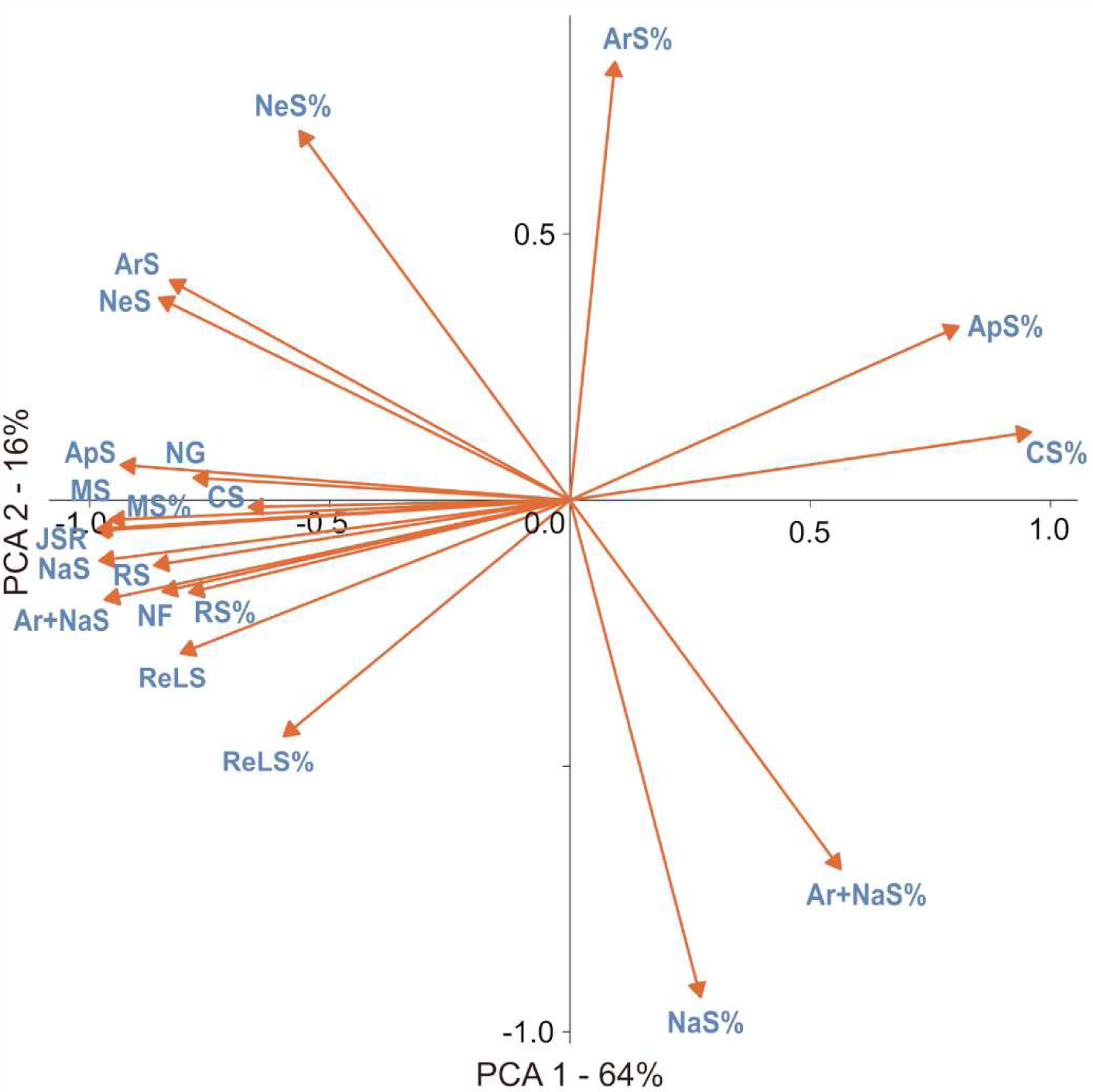
Results of PCA ordination. For simplification, only the first two PCA axes are shown.

**Table 2.** **P**rincipal component analysis (PCA) results, including correlations between the variable and PCA axis and the percentage variation explained by particular axis. Correlations with an absolute value larger than 0.5 are highlighted in bold.

| Variable | PCA 1 | PCA 2 | PCA 3 |
| --- | --- | --- | --- |
| JSR = Joined species richness | **-0.99** | -0.06 | 0.07 |
| NaS = Native species richness | **-0.97** | -0.19 | 0.09 |
| NaS_ '%' = ( NaS /JSR) × 100 | 0.27 | **-0.93** | 0.11 |
| ReLS = Red list species | **-0.81** | -0.29 | -0.40 |
| ReLS_ '%' = (ReLS/JSR) × 100 | **-0.60** | **-0.44** | **-0.51** |
| NeS = Neophyte species | **-0.86** | 0.38 | -0.18 |
| NeS_ '%' = (NeS/JSR) × 100 | **-0.56** | **0.69** | -0.23 |
| ArS = Archeophyte species | **-0.83** | **0.41** | 0.12 |
| ArS_ '%' = ( ArS /JSR) × 100 | 0.09 | **0.82** | 0.04 |
| 'Ar+Na'S = Archeophyte and native species | **-0.98** | -0.11 | 0.10 |
| 'Ar+Na'S_ '%' = ( 'Ar+Na'S /JSR) × 100 | **0.56** | **-0.69** | 0.23 |
| ApS = Apophyte species | **-0.93** | 0.07 | 0.17 |
| ApS_ '%' = ( ApS /JSR) × 100 | **0.81** | 0.33 | -0.07 |
| RS = Rare species | **-0.87** | -0.12 | **-0.42** |
| RS_ '%' = ( RS /JSR) × 100 | **-0.79** | -0.17 | **-0.52** |
| MS = Moderate species | **-0.96** | -0.04 | 0.23 |
| MS_ '%' = ( MS/JSR) × 100 | **-0.67** | -0.01 | **0.65** |
| CS = Common species | **-0.78** | 0.04 | **0.49** |
| CS_ '%'=(CS/JSR)*100 | **0.96** | 0.13 | -0.04 |
| NG= Number of genera | **-0.98** | -0.05 | 0.12 |
| NF= Number of family | **-0.85** | -0.17 | 0.11 |
| eigenvalue | 13.5 | 3.4 | 1.8 |
| variance.percent | 64.3 | 16.0 | 8.7 |
| cumulative.variance.percent | 64.3 | 80.3 | 89.1 |

The native species hot spots included squares with at least 636 native species present. They were rather clustered, creating 52 separate areas, and three main large areas can be distinguished: (1) at the junction of Lublin and Kielce uplands, (2) in the Kielce and Przedbórz uplands, and (3) in the Silesia and Kraków–Częstochowa Uplands together with the Oświęcim Basin. Two squares with the highest richness of native species were in the Wisła River valley in Puławy (square FE13 with 876 species) and also in the Wisła River valley in the eastern and central part of Kraków (DF69, 874 species). The native species cold spots included squares with less than 216 species, strongly dispersed into 71 squares in separate areas. The largest of these areas was in the Northern Masovia Lowland.

The location of archeophyte hot spots (squares with at least 92 archeophyte species recorded) overlapped to a large extent with native SR and were concentrated in the Lublin and Kielce uplands and also included (1) the eastern part of Southern Masovia Hills, (2) the Nida Basin and Kielce and Przedbórz uplands, and (3) Silesia Upland. The two most species-rich squares for archeophytes were in Silesia, including the northern part of Ople city (square CE95 with 131 species) and the northeastern part of Wrocław (BE49, 128 species). They were rather dispersed over 60 separate areas. The archeophyte cold spots included squares with less than 25 species and were dispersed in 62 areas. The largest of theses areas were in Northern Podlasie Plain and Masurian Lakeland.

The largest concentration of the red list species hot spots (squares with at least 76 species) were found (1) in Western and Vohlyn Polesie, (2) again at the junction of Lublin and Kielce Uplands, and (3) at the junction of Lower Vistula River valley and Toruń-Eberswalde ice-marginal valley. However, the dispersion of red list species hot spots was higher than that of native and archeophytes, with the inclusion of numerous smaller areas such as the Kraków-Częstochowa uplands, Roztocze upland, and areas near the Pozna and Wielkopolski National Park in Wielkopolskie Lakeland. Two squares that were the most species rich for red list species included Gdańsk (square DA80, with 181 species) and the northwestern part of Kraków (square DF69 173). They were rather clustered, forming 51 separate areas. The red list cold spots included squares with less than six species, grouped in 60 areas. The largest of these areas were in Northern Masovia Lowland and Southern Wielkopolska Lowland.

With regard to neophyte hot spots (squares with 65 or more neophyte species), strong concentrations were apparent in Silesia Upland and Oświęcim Basin, and also near Szczecin and Poznań, and quite surprisingly, in Sandomierz Basin between Przeworsk and Rzeszów, among other areas. The two squares that were the most rich in neophytes included the northeastern part of Wrocław (BE 49, 155 species) and Poznań centre (BD08, 134). The hot spots were dispersed in 60 separate areas. The neophyte cold spots included squares with less than seven neophyte species confirmed. The cold spots were highly concentrated in 36 areas, with the largest ones located in Northern Podlasie Plain and Northern Masovia Lowland.

Unfortunately, the location of native, archeophytes, and red list species did not overlap with the presence of national parks (Fig 4a). Moreover, the native and archeophyte species-rich areas coincided with neophyte hot spots (4b).

## 4. Discussion

The main patterns of SR and the percentages in a particular group can be attributed to three components: overall SR (PCA 1), invasion level (PCA 2), and flora peculiarity of some squares resulting from high number and fraction of rare species, often with a high conservation value (PCA 3). It should be highlighted that some correlations (e.g., between native SR and red list species) resulted not only from causal reasons but were also statistical; for example, the native species group included numerous red list species. The distribution of species range sizes in Poland, expressed as the number of occupied squares, did not differ from those typically observed, as most of the species have a limited range (Lennon et al., 2004). Although the rare species constituted a major part of the species pool in a region, spatial structures of SR were typically dominated by the more common species (Lennon et al., 2004). Here, we also observed that the SR of species with a moderate distribution range (that is, moderately frequent species) was strongly correlated with joined SR (**Table 2** and **Table S2**). The results underline that preserving biodiversity in Poland will require focusing not only on rare or red list species, but also having landscapes that are able to maintain numerous species with moderate distribution. The presence of such species is crucial to biodiversity protection, especially outside mountainous areas (**Fig 3f**). The protection of entire landscapes with a high fraction of semi-natural habitats seems to be especially important in the face of anthropogenic climate and land use changes. For example, a large-scale survey in Great Britain revealed that areas with a high fraction of semi-natural habitats (e.g., grasslands, wetlands heartlands) are relatively resistant to negative changes of biodiversity (Monteràs-Janera et al., 2022). Moreover, the results obtained suggest that, at 10 × 10 km spatial resolution, a high percentage of apophytes in a landscape does not necessarily result from high anthropogenic pressure, but could be a statistical effect of overall low SR (**Tab 2, Fig 4**) owing to a low number of rare and moderately frequent species.

A visible overall trend of SR decrease was apparent from the southwest towards the northeast in Poland. This trend generally corresponds with the continental pattern of vascular plant SR in Europe (Mutke & Barthlott, 2005; Araújo et al., 2005; Cai et al., 2022; **Fig 1a**). However, beyond the large-scale variation in SR, local factors such as soil pH shape SR (Pärtel et al., 2016) and thus disrupt the broader pattern in specific locations. For example, the present-day regional diversity of archeophytes mirrors the intensity of past human settlements (Macek et al., 2022). In Poland, recent low settlement density may be a continuation of low interest in settlement in certain areas in the past (Karpińska-Kołaczek et al., 2014). This hypothesis is suggested by studies from the Czech Republic that confirmed repeated settlement over millennia in the same ecological regions (Dejmán et al., 2022). In the case of Poland, settlement was strongly related to ice sheet extension during the Last Glaciation Maximum and the development of loess sediment in ice-free areas (**Fig 1a**). The areas in the south of Poland were thus preferentially inhabited in the Paleolithic Age (Moskal-del Hoyo et al., 2021). Another factor, overriding the biogeographical pattern resulting in a north-south gradient in SR in Europe, is the land relief of Poland. The majority of the country’s territory is relatively flat and characterized by lowlands, while the mountains and highlands are located in the south. As a result, the mountain species, including the majority of endemic species of Poland (Mirek & Pięknoś-Mirkowa, 2009), are absent in the central and northern parts of Poland. Consequently, the areas with high conservation value are often in southern Poland, revealing similar trends to overall SR (**Fig. 5d**). Another consequence of this land-relief peculiarity of Poland is difficulty linking observed SR with the distance from the south European glacial refugia. The pan-European gradient of decreasing SR from south to north is correlated, among other factors, with constrained species dispersal from glacial refugia (Svenning et al., 2007, 2008; Normand et al., 2011), and it is tempting to exploit this explanation to explain the SR pattern in Poland. However, in our opinion, on a country-level scale, the effect of land relief and/or geology of Poland overrides the effect of dispersal limitation, especially considering the shorter distance range in Poland compared with the entire European continent.

**Figure 5.**
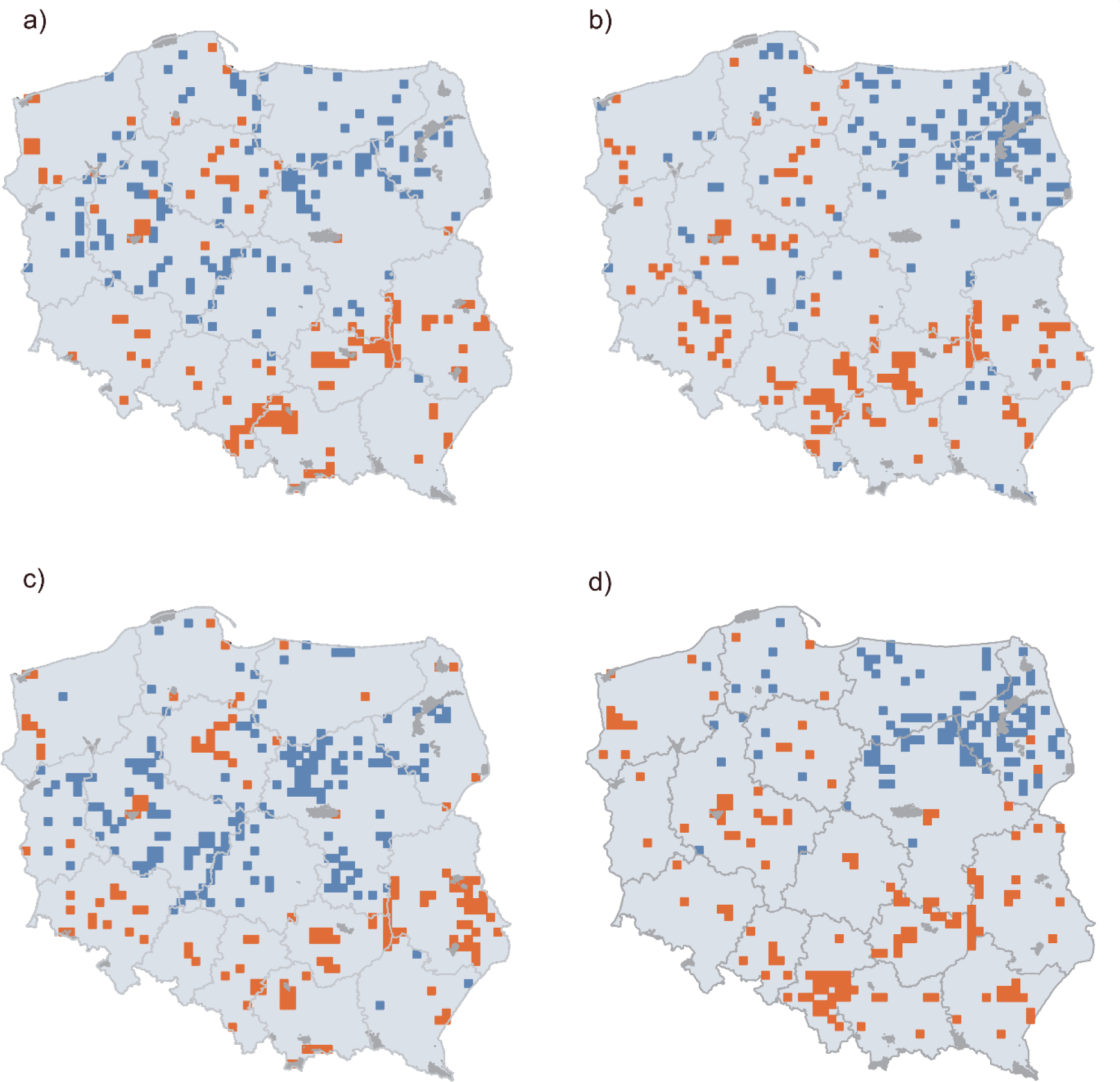
Location of squares identified as hot spots (red) and cold spots (blue) of native (a), archeophytes (b), red list (c), and neophytes (d) on the background of voivodeship boundaries (dark grey lines). The locations of national parks are also shown (dark grey).

The high native SR and presence of SR hot spots are, unfortunately, also positively correlated with the presence of neophytes (**Fig. 4, Table S2**). The positive correlations between native and invasive SR were always observed at larger spatial extents (Peng et al., 2019). This finding can be explained by “spatial heterogeneity” and “favorable conditions” hypotheses. The spatial heterogeneity hypothesis assumes that at larger spatial scales, there is likely to be greater resource heterogeneity, allowing more species to coexist (Hawkins et al., 2003). The favorable conditions hypothesis predicts that in overall favorable conditions (e.g., with available resources), more species can coexist (de Albuquerque et al., 2011).

### 4.1. Hotspot protection and invasion risks

The results reveal that numerous hot spots of vascular plant SR in Poland—including areas in Kielce Upland, Oświęcim Basin, Kraków-Częstowchow Upland, Lower Vistula River valley, Toruń-Ebersvald ice-marginal valley, and Szczecin Coastland—are not protected as national parks. The new projects raised by non-government organizations (Klub, 2023) can, to some extent, help by proposing that national parks be established in areas such as Puszcza Bukowa (Szczecin Coastland), Jurajski National Parks (Kraków-Częstochowa Upland), and Odrzański National Park (Silesia Lowland). Additionally, as mentioned previously, the problem of correlation of native SR with neophytes richness, and in practice, invasive alien species are a major problem for managers of protected areas in Central Europe (Braun et al., 2016) and worldwide (Foxcroft et al., 2017).

### 4.2. Data set completeness and temporal adequateness

Data regarding SR in particular regions are often incomplete. To solve this bias, different data sources are combined, but harmonization of taxonomical nomenclature is needed (Wüest et al., 2020; Grenié et al., 2023). Since plant nomenclature is not consistent across time, informal species aggregations are often created to acknowledge historical data and so-called microspecies that are hard to identify precisely (Dengler et al., 2012; Pescott et al., 2018). The total number of taxa examined in the current study was 2137, while the recent checklist of vascular plant species in Poland denoted 2600 native species and 534 neophytes (Mirek et al., 2020). Thus, our maps contain information on about 69% of all species. However, the function of a checklist is to include the largest number of well-defined taxa (Mirek et al., 2020), while for SR mapping, realistic data on distribution are necessary. For example, knowledge regarding the distribution of 550 species representing merely four genera (*Taraxacum*, *Rubus*, *Hieracium*, and *Alchemilla*) is limited to restricted areas explored by specialists focusing on a given genera. In practice, a similar fraction of species representing national flora is possible to analyze; for example, the analysis of SR changes of vascular plants in Germany relies on a comparison of 77% of the entire country’s flora (Eichenberg et al., 2021), while in the Czech Republic, a similar analysis is based on 54% of species listed on the Czech checklist (Klinkovská et al., 2024). Unfortunately, the projects from which the SR data were derived were not designed to track species distribution in time.

Given the main effort for ATPOL project realization and the years of plot recording in PVD, the observed patterns of SR seem to mostly reflect the situation at the end of the 20th century; nonetheless, the data regarding neophytes were updated in 2019 (Zając & Zając, 2019). In a comparison of data from Germany (Eichenberg et al., 2020), loss of about 3% of native species and 5.3% of archeophytes in average SR per 5 × 5 km grid cells occurred between the periods of 1988-1996 and 1997-2017. Data from the Czech Republic (Klinkovská et al., 2024) showed a diminishing range (number of squares with presence) of specialists in nutrient-poor, frequently but not-intense disturbed habitats, with low colonization and competitive abilities. Many of the species were included in the national red list. The decrease was linked to an ongoing decline in habitat quality after the cessation of traditional management (Klinkovská et al., 2024), and a similar trend could be assumed to be occurring in Poland as well. The decrease of native and archeophytes SR in Poland is most likely not spatially homogenous; for example, the decrease of arable weeds SR was observed in lowlands with intense agriculture, but not in mountains and highlands with more extensive land use (Dąkowska et al., 2017). Similarly, results of the comparison of repeated vegetation surveys reveal different patterns of species composition and SR changes regarding studied vegetation (e.g., rocky outcrops vs forests; Reczyńska & Świerkosz, 2017, 2024).

### 4.3. Potential sampling bias

Typically, higher sampling effort occurs in areas with easy access and that are interesting for researchers (e.g., Szymura et al., 2023a). In the case of our data set, some doubts can be related to observed low SR in the northwestern part of Poland and high SR in cities with old universities (e.g., Wrocław, Kraków, and Poznań). The northwestern part of Poland could be undersampled owing to the relatively large distance from academic centers and the low density of urbanized areas and communication routes. However, the completeness of records from this part of Poland was checked by planned field campaigns of trained botanists, and the results generally confirmed low SR (A. Zając, personal communication). Moreover, potentially well-sampled areas such as Biebrzański, Białowieża, and Wigierski National Parks were not considered as hot spots, and in Biebrzański National Parka a cold spots are located. Finally, at the pan-European scale, the northeastern region of Poland is recognized as being species-poor, which results from the continental gradient of decreasing SR toward the northeast (e.g., Cai et al., 2022, **Fig 1b**). The low SR is only poorly enhanced by archeophytes due to a lack of loess sediments (Lehmkuhl et al., 2021) and more fertile soils, which correlates with settlement location during the Neolithic Revolution (Nowak, 2013; Moskal-del Hoyo et al., 2021). Low sampling effort is not expected in the mountains since, in the contemporary territory of Poland, mountainous areas were traditionally explored by botanists (for discussion see Szymura et al., 2023a). Regarding the high SR found in cities, the European cities’ flora is naturally species-rich (Kühn et al., 2004), and globally, cities are often located in areas of high biodiversity; thus, urbanization is relatively higher in areas with high biodiversity (Kühn et al., 2004; Luck, 2007; Ives et al., 2016). The broad-scale positive correlation between human presence and SR suggests that people have preferentially settled and generally flourished in areas of high biodiversity and/or have contributed to it with species introductions and habitat diversification (Pautasso, 2007). The large, old towns (e.g., Wrocław, Karków, and Poznań) were settled in areas with well-developed agriculture, which can explain the high number of archeophytes. Moreover, it should be highlighted that the SR is calculated for squares 10 × 10 km that include not only strict city centers but also suburban and rural areas; in addition, the data also contained historical records from times with relatively low urbanization and agriculture intensity. The pattern of richness of neophytes, which are the most numerous in cities and in densely populated and highly industrialized areas of the Silesia upland, seems to be very likely considering neophyte introduction and pathways of further spread (Kovarik & Lippe, 2007; Tokarska-Guzik et al., 2008). To conclude, we argue that although some bias towards undersampling of less-accessible areas holding little interest for botanists and high sampling effort near cities with universities is likely, the general pattern observed here results predominantly from ecological processes.

## Conclusions

- The richness of native species, archeophytes, neophytes, and red list species is positively correlated in Poland
- The main patterns of species richness patterns in Poland can be interpreted as (1) a general gradient of species richness, (2) differentiation in the number and percentage of neophytes, and (3) gradient in the number and percentage of red list and rare species in local flora.
- In general, the northeastern part of Poland is species-poor, while the highest species richness is concentrated in the mountains and uplands of southern Poland and in some parts of the Wisła River valley.
- The recent system of national parks in Poland does not overlap with national-scale hot spots of native, archeophytes, and red list species.
- The correlation between native species and neophytes richness indicates that biological invasions are a serious threat to biodiversity maintenance in Poland.
- Presented maps seem to reflect general ecological gradients influencing species richness in Poland, in addition to likely spatial and temporal biases.

## Financing

This work was supported by the National Science Centre, Poland, project number 2019/35/B/NZ8/00273.

## Authors’s contributions

**THS** – Conceptualization, Data curation, Formal analysis, Methodology, Funding acquisition, Project administration, Writing the manuscript. **HT** – Data curation, Formal analysis, Writing the manuscript. **MS**-Conceptualization, Methodology, Writing the manuscript

## Appendix

**Fig S1.**
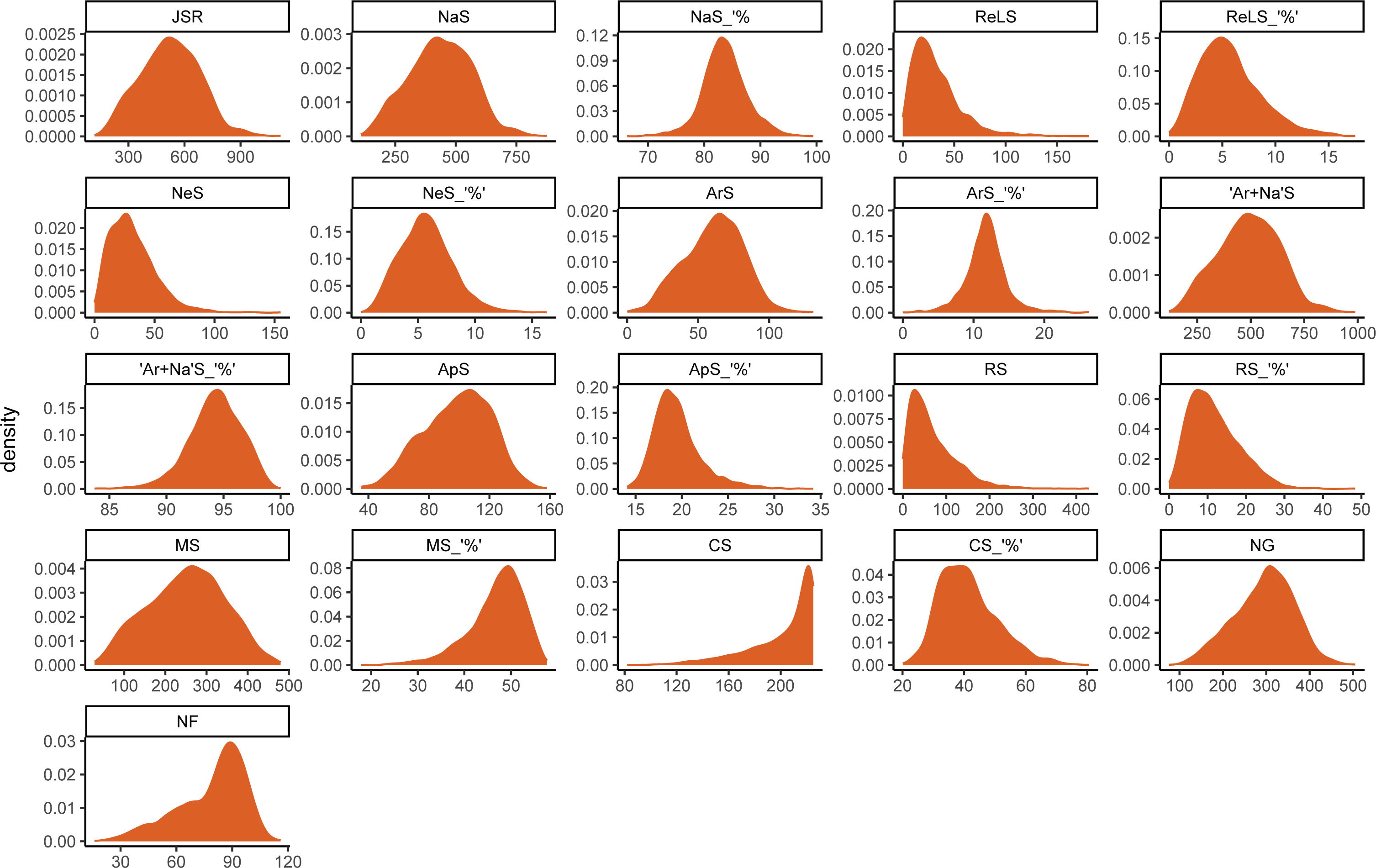
Distribution of species richness in particular species groups in squares.

**Table S1.** Descriptive statistics of species richness in species groups.

|  | median | mean | std.dev | coef.var [%] | min | max | range |
| --- | --- | --- | --- | --- | --- | --- | --- |
| **JSR= Joined species richness** | 522 | 522.3 | 158.3 | 0.30 | 119 | 1114 | 995 |
| **NaS= Native species richness** | 437 | 435.6 | 128.8 | 0.30 | 108 | 876 | 768 |
| **NaS_ '%'=( NaS /JSR)\*100** | 83.5 | 83.7 | 3.9 | 0.05 | 66.3 | 99.4 | 33.1 |
| **ReLS= Red list species** | 27 | 32.3 | 23.6 | 0.73 | 0 | 181 | 181 |
| **ReLS_ '%'=(ReLS/JSR)\*100** | 5.3 | 5.7 | 2.9 | 0.50 | 0 | 17.5 | 17.5 |
| **NeS= Neophyte species** | 29 | 31.5 | 18.8 | 0.60 | 0 | 155 | 155 |
| **NeS_ '%'=( NeS /JSR)\*100** | 5.6 | 5.7 | 2.3 | 0.40 | 0 | 16.3 | 16.3 |
| **ArS= Archeophyte species** | 62 | 60.9 | 20.5 | 0.34 | 0 | 131 | 131 |
| **ArS_ '%'=( ArS /JSR)\*100** | 11.8 | 11.8 | 2.8 | 0.24 | 0 | 26.3 | 26.3 |
| **'Ar+Na'S= Archeophyte and native species** | 493 | 490.7 | 143.3 | 0.29 | 119 | 988 | 869 |
| **'Ar+Na'S_ '%'=( 'Ar+Na'S /JSR)\*100** | 94.4 | 94.3 | 2.3 | 0.02 | 83.7 | 100 | 16.3 |
| **ApS= Apophyte species** | 100 | 98.6 | 22.0 | 0.22 | 35 | 158 | 123 |
| **ApS_ '%'=( ApS /JSR)\*100** | 19.1 | 19.5 | 2.6 | 0.13 | 14.1 | 34.2 | 20.1 |
| **RS= Rare species** | 53 | 69.0 | 55.7 | 0.81 | 0 | 429 | 429 |
| **RS_ '%'=( RS /JSR)\*100** | 10.5 | 11.7 | 6.5 | 0.55 | 0 | 48.4 | 48.4 |
| **MS= Moderate species** | 256 | 251.2 | 92.2 | 0.37 | 28 | 480 | 452 |
| **MS_ '%'=( MS/JSR)\*100** | 47.9 | 46.9 | 5.8 | 0.12 | 17.8 | 57.7 | 39.9 |
| **CS= Common species** | 212 | 202.1 | 25.5 | 0.13 | 82 | 225 | 143 |
| **CS_ '%'=(CS/JSR)\*100** | 40 | 41.4 | 9.4 | 0.23 | 20.1 | 80.5 | 60.4 |
| **NG= Number of genera** | 301 | 294.9 | 68.6 | 0.23 | 76 | 505 | 429 |
| **NF= Number of family** | 84 | 78.9 | 18.0 | 0.23 | 16 | 116 | 100 |

**Table S2.**
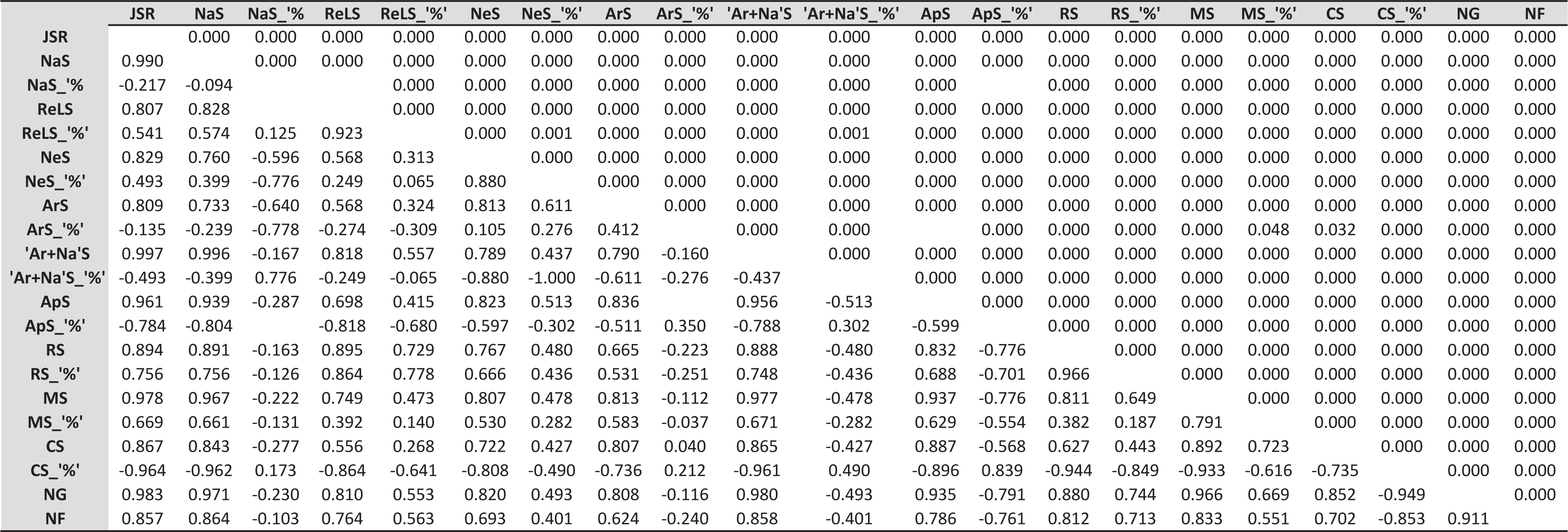
Spearman rank correlations between the richness of species in different groups (Table S2). The upper matrix triangle shows p values, while the lower shows r. Nonsignificant correlations are not shown.

|  | JSR | NaS | NaS_ '%' | ReLS | ReLS_ '%' | NeS | NeS_ '%' | ArS | ArS_ '%' | 'Ar+Na'S | 'Ar+Na'S_ '%' | ApS | ApS_ '%' | RS | RS_ '%' | MS | MS_ '%' | CS | CS_ '%' | NG | NF |
| --- | --- | --- | --- | --- | --- | --- | --- | --- | --- | --- | --- | --- | --- | --- | --- | --- | --- | --- | --- | --- | --- |
| JSR |  | 0.000 | 0.000 | 0.000 | 0.000 | 0.000 | 0.000 | 0.000 | 0.000 | 0.000 | 0.000 | 0.000 | 0.000 | 0.000 | 0.000 | 0.000 | 0.000 | 0.000 | 0.000 | 0.000 | 0.000 |
| NaS | 0.990 |  | 0.000 | 0.000 | 0.000 | 0.000 | 0.000 | 0.000 | 0.000 | 0.000 | 0.000 | 0.000 | 0.000 | 0.000 | 0.000 | 0.000 | 0.000 | 0.000 | 0.000 | 0.000 | 0.000 |
| NaS_ '%' | -0.217 | -0.094 |  |  | 0.000 | 0.000 | 0.000 | 0.000 | 0.000 | 0.000 | 0.000 | 0.000 |  | 0.000 | 0.000 | 0.000 | 0.000 | 0.000 | 0.000 | 0.000 | 0.000 |
| ReLS | 0.807 | 0.828 |  |  | 0.000 | 0.000 | 0.000 | 0.000 | 0.000 | 0.000 | 0.000 | 0.000 | 0.000 | 0.000 | 0.000 | 0.000 | 0.000 | 0.000 | 0.000 | 0.000 | 0.000 |
| ReLS_ '%' | 0.541 | 0.574 | 0.125 | 0.923 |  | 0.000 | 0.001 | 0.000 | 0.000 | 0.000 | 0.001 | 0.000 | 0.000 | 0.000 | 0.000 | 0.000 | 0.000 | 0.000 | 0.000 | 0.000 | 0.000 |
| NeS | 0.829 | 0.760 | -0.596 | 0.568 | 0.313 |  | 0.000 | 0.000 | 0.000 | 0.000 | 0.000 | 0.000 | 0.000 | 0.000 | 0.000 | 0.000 | 0.000 | 0.000 | 0.000 | 0.000 | 0.000 |
| NeS_ '%' | 0.493 | 0.399 | -0.776 | 0.249 | 0.065 | 0.880 |  | 0.000 | 0.000 | 0.000 | 0.000 | 0.000 | 0.000 | 0.000 | 0.000 | 0.000 | 0.000 | 0.000 | 0.000 | 0.000 | 0.000 |
| ArS | 0.809 | 0.733 | -0.640 | 0.568 | 0.324 | 0.813 | 0.611 |  | 0.000 | 0.000 | 0.000 | 0.000 | 0.000 | 0.000 | 0.000 | 0.000 | 0.000 | 0.000 | 0.000 | 0.000 | 0.000 |
| ArS_ '%' | -0.135 | -0.239 | -0.778 | -0.274 | -0.309 | 0.105 | 0.276 | 0.412 |  | 0.000 | 0.000 |  | 0.000 | 0.000 | 0.000 | 0.000 | 0.048 | 0.032 | 0.000 | 0.000 | 0.000 |
| 'Ar+Na'S | 0.997 | 0.996 | -0.167 | 0.818 | 0.557 | 0.789 | 0.437 | 0.790 | -0.160 |  | 0.000 | 0.000 | 0.000 | 0.000 | 0.000 | 0.000 | 0.000 | 0.000 | 0.000 | 0.000 | 0.000 |
| 'Ar+Na'S_ '%' | -0.493 | -0.399 | 0.776 | -0.249 | -0.065 | -0.880 | -1.000 | -0.611 | -0.276 | -0.437 |  | 0.000 | 0.000 | 0.000 | 0.000 | 0.000 | 0.000 | 0.000 | 0.000 | 0.000 | 0.000 |
| ApS | 0.961 | 0.939 | -0.287 | 0.698 | 0.415 | 0.823 | 0.513 | 0.836 |  | 0.956 | -0.513 |  | 0.000 | 0.000 | 0.000 | 0.000 | 0.000 | 0.000 | 0.000 | 0.000 | 0.000 |
| ApS_ '%' | -0.784 | -0.804 |  | -0.818 | -0.680 | -0.597 | -0.302 | -0.511 | 0.350 | -0.788 | 0.302 | -0.599 |  | 0.000 | 0.000 | 0.000 | 0.000 | 0.000 | 0.000 | 0.000 | 0.000 |
| RS | 0.894 | 0.891 | -0.163 | 0.895 | 0.729 | 0.767 | 0.480 | 0.665 | -0.223 | 0.888 | -0.480 | 0.832 | -0.776 |  | 0.000 | 0.000 | 0.000 | 0.000 | 0.000 | 0.000 | 0.000 |
| RS_ '%' | 0.756 | 0.756 | -0.126 | 0.864 | 0.778 | 0.666 | 0.436 | 0.531 | -0.251 | 0.748 | -0.436 | 0.688 | -0.701 | 0.966 |  | 0.000 | 0.000 | 0.000 | 0.000 | 0.000 | 0.000 |
| MS | 0.978 | 0.967 | -0.222 | 0.749 | 0.473 | 0.807 | 0.478 | 0.813 | -0.112 | 0.977 | -0.478 | 0.937 | -0.776 | 0.811 | 0.649 |  | 0.000 | 0.000 | 0.000 | 0.000 | 0.000 |
| MS_ '%' | 0.669 | 0.661 | -0.131 | 0.392 | 0.140 | 0.530 | 0.282 | 0.583 | -0.037 | 0.671 | -0.282 | 0.629 | -0.554 | 0.382 | 0.187 | 0.791 |  | 0.000 | 0.000 | 0.000 | 0.000 |
| CS | 0.867 | 0.843 | -0.277 | 0.556 | 0.268 | 0.722 | 0.427 | 0.807 | 0.040 | 0.865 | -0.427 | 0.887 | -0.568 | 0.627 | 0.443 | 0.892 | 0.723 |  | 0.000 | 0.000 | 0.000 |
| CS_ '%' | -0.964 | -0.962 | 0.173 | -0.864 | -0.641 | -0.808 | -0.490 | -0.736 | 0.212 | -0.961 | 0.490 | -0.896 | 0.839 | -0.944 | -0.849 | -0.933 | -0.616 | -0.735 |  | 0.000 | 0.000 |
| NG | 0.983 | 0.971 | -0.230 | 0.810 | 0.553 | 0.820 | 0.493 | 0.808 | -0.116 | 0.980 | -0.493 | 0.935 | -0.791 | 0.880 | 0.744 | 0.966 | 0.669 | 0.852 | -0.949 |  | 0.000 |
| NF | 0.857 | 0.864 | -0.103 | 0.764 | 0.563 | 0.693 | 0.401 | 0.624 | -0.240 | 0.858 | -0.401 | 0.786 | -0.761 | 0.812 | 0.713 | 0.833 | 0.551 | 0.702 | -0.853 | 0.911 |  |

